# Empowering multitrait brain phenotypes GWAS

**DOI:** 10.1101/2025.05.04.652120

**Authors:** Auvergne Antoine, Nicolas Traut, Léo Henches, Hanna Julienne, Arthur Frouin, Roberto Toro, Hugues Aschard

**Affiliations:** Institut Pasteur, Université Paris Cité, Department of Computational Biology, F-75015 Paris, France; Department of Epidemiology, Harvard T. H. Chan School of Public Health, Boston, Massachusetts, United States

## Abstract

Brain magnetic resonance imaging (MRI) data are now becoming available in human genetic cohorts including thousands of participants, providing a powerful mean to decipher the genetic architecture of brain phenotypes. Univariates genome-wide association screenings (GWAS) are typically applied as a baseline approach, but there are increasing interest for multitrait approaches to both increase statistical power and uncover shared structure underlying those phenotypes. However, there are limited guidelines on how to select phenotypes to be analyzed jointly and optimize power. Here we investigated factors impacting variant discoverability of state-of-the art multitrait GWAS using 1,010 MRI-derived phenotypes measured in 32,947 participants from the UK Biobank. Our study shows that the expected gain of multitrait approaches over univariate GWAS is highly predictable. We further propose data-driven strategy to define optimize sets and compare its performances with biologically driven approaches.

## Main

One of the first significant relationship identified between a brain phenotype and a psychiatric disorder in 1976^1^ raised the interest of researchers for quantitative biomarkers, and led to the emergence of research programs investigating links between psychotherapy and neuroimaging^2–4^. Since then, systematic brain mapping atlases have been developed^5^ to help standardizing and organizing the increasing amount of neuroimaging data available, and many noninvasive techniques have been developed, including structural magnetic resonance imaging (MRI), functional MRI (fMRI), and magnetic resonance spectroscopy (MRS). Among the various approaches developed to study brain characteristics, neuroimaging genetics has gained considerable momentum as a mean to decipher the genetic pathway underlying the human brain, pathways that may ultimately be mapped to psychiatric disorder. However, with modern cohorts including thousands of MRI-derived phenotypes and millions of genetics variants measured in thousands of individuals, achieving this goal faces both combinatorial and computational challenges.

Following standard practice in polygenic phenotypes study, past MRI research typically conducted systematic univariate genome-wide association studies (GWAS), examining association between each genotype and a single phenotype at a time as primary analyses^6–9^. Given the dimensionality of existing data and the limited statistical power of GWAS to detect variants with modest effect, an increasing number of studies are applying multitrait GWAS to boost variants identification^8–12^. Briefly, a multitrait GWAS consists in testing jointly the effect of each variant with a set of phenotypes in a single test (using a composite null hypothesis H0 being: *β*_1_ = *β*_2_ = … *β*_*K*_ = 0, where *β*_*k*=l…*K*_ are the effect of the variant test on phenotype *k*). Using such method has proven useful in both neuroimaging genetics and in the study of other traits^13^ but raise the question of which phenotype to analyze jointly. This is not a trivial question, and whatever the sets considered, they will only represent a tiny fraction of all possibilities (for example, for 1,000 phenotypes, there are over 10^300^ possible combinations of phenotypes).

Here we examined the factors impacting the performances of multitrait brain imaging GWAS, assess the relative power of knowledge-driven selection strategies, and ultimately devise a data-driven approach to optimize trait’s set selection. We used 1,010 brain phenotypes and 9,611,512 variants measured in 33,006 unrelated European participants from the UK Biobank. The phenotypes include whole brain/left and right hemisphere phenotypes: volumes, area, thickness, grey/white matter contrast, mean intensity and number of holes (**Table S1**). Univariate GWAS was conducted using PLINK 2.0 for each phenotype and used as a baseline. We then sampled hundreds of sets of phenotypes based on various criteria and assessed the gain of multitrait GWAS over univariate GWAS as the ratio of independent loci detected by either approach, referred further as *τ*_*gain*_. Importantly, conducting such an analysis directly from the individual-level data would carry a tremendous computational cost (**supplementary Notes**). To circumvent this issue, we used the JASS framework^13,14^ where all multitrait GWAS analyses are derived from the univariate GWAS summary statistics, considerably reducing the computational burden. Guidelines on how to pre-process the GWAS summary statistics, quality control, and the derivation of the necessary complementary information for summary statistics-based multitrait analyses are provided in the **Online Methods** and **Supplementary Material** (**Notes 1** and **Fig. S1-2**). The JASS framework can accommodate various multivariate statistics. We used an omnibus test, a model that has been shown to provide a good tradeoff in power across a range of scenarios^15^, and that is equivalent to a standard two-way MANOVA in individual-level data.

We first conducted an agnostic experiment, sampling 2750 sets of size *K* = 5 to 200 phenotypes from the entire pool of 1,010 phenotypes. For each set we measured *τ*_*gain*_, the detection gain for the multitrait test for each set, and examined its correlation with seven parameters: the set size, the average of trait’s heritability, the average trait’s polygenicity, the mean absolute genetic and phenotypic correlations, the number of significant loci in the univariate analyses, and the condition number of the phenotypic correlation, which quantifies the amount of collinearity in the set (see **Online Methods**). The average *τ*_*gain*_ across all set equals 1.117 Overall, we observed a 12% increase in detection for the multitrait test. The gain increases with the size of the sets, with the average gain rising from 0.997 for set size of 5, to 1.328 for set size of 200 phenotypes (**Fig. 1a**). All seven parameters, except the average genetic correlation show strong and positive marginal association with the gain (**Table S2**). Taken together in a joint model those features show a highly significant predictive power of the gain with an adjusted R-squared of 0.30 (*P* = 2.4e-204). A sensitivity analysis using a linear mixed model to account for the overlap between the sets of phenotypes produced qualitatively similar results (**Table S2, Online Methods**). Given the large effect of set size, we further considered a stratified model, estimating the effects of the six remaining parameters for each set size. These set size-specific models displayed much higher fit, with R-squared ranging from 0.65 (P = 5.9 × 10^−53^, size = 40) to 0.77 (P = 6.9 × 10^−76^, size = 15). The parameter’s estimates varied substantially with set size (**Fig. 1b-c, Table S3**). The average heritability and the number of significant loci in the univariate analyses become less important with increasing set size, and conversely the conditional number moves from being non-significant for small set to become the primary predictor of the gain for large set.

**Figure 1.**
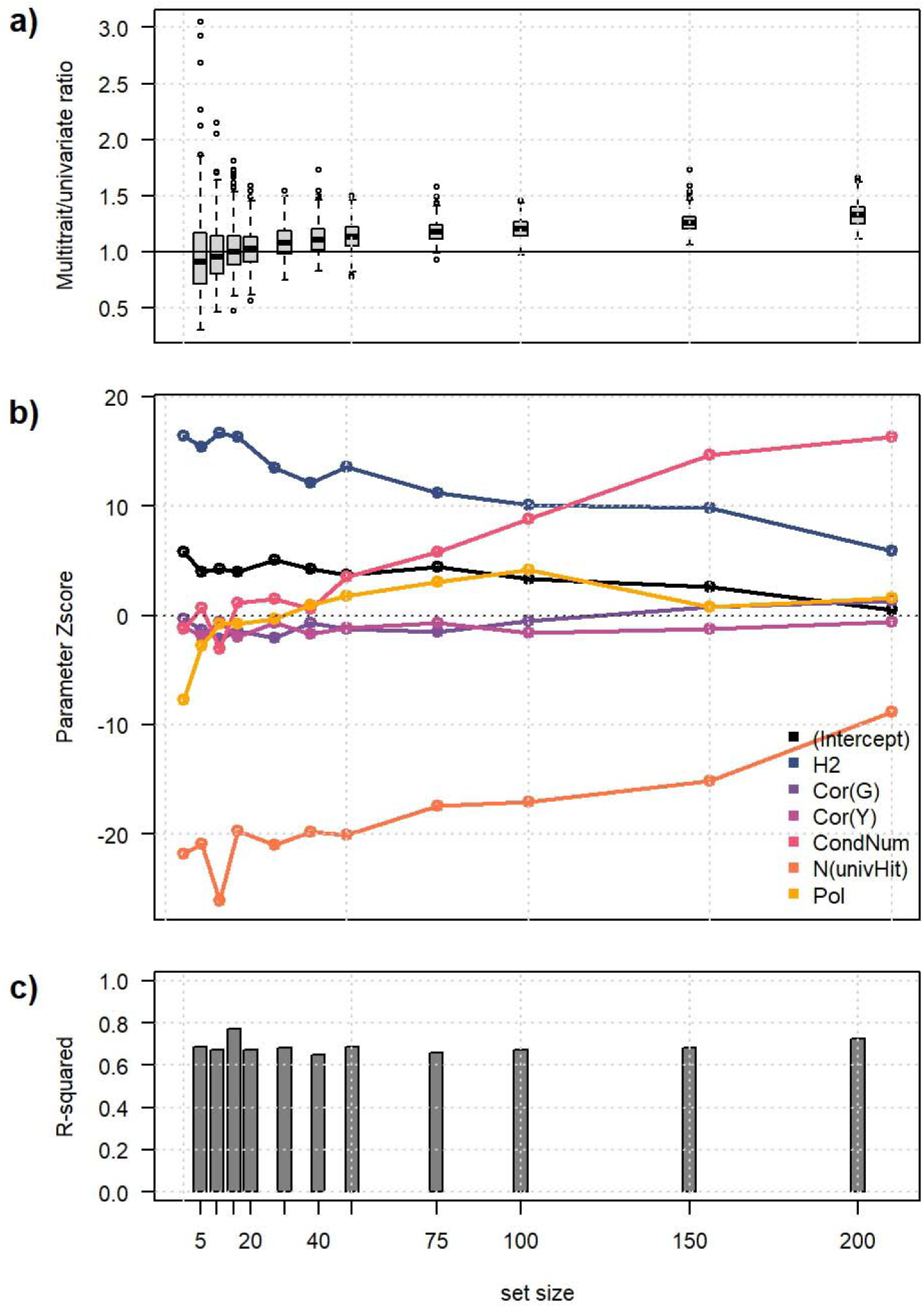
Multitrait GWAS gain and its main driver. We sampled sets of 5 to 200 phenotypes from a total of 1,010 phenotypes. **(a)** For each set we derived the multitrait power gain, *τ*_*gain*_, as the ratio of independent loci detected by the multitrait test as compared to the univariate test. **(b)** For each set size considered, we fitted a joint model estimating the effect of six parameters: the average phenotype heritability (H2) and polygenicity (Pol), the number of univariate hits identified (N(univHit)), the phenotypic (Cor(Y)) and genetic (Cor(G)) correlation, and the condition number of the phenotypic correlation matrix (CondNum). **(c)** For each set size considered, we derived the adjusted R squared of the joint model.

We examined the behavior of the predictive model across sets defined based on trait relatedness, an approach commonly used in multitrait screening^8,9^. We considered eight groups described in **Table S4** and referred as *area, volume, intensity, basal, neocortex, thickness, contrast*, and *limbic*. The predicted values show strong correlation with the observed gain overall and within each group (**Fig. 2a, Fig. S3**). This confirm that in many scenarios, the expected gain from multitrait analysis can be derived based on derived traits’ characteristics, independently of the known trait’s biological relatedness. Although, as showed in the derived mean squared error (MSE) from **Figure 2c** and the detailed plots from **Figures S3**, three groups display a partial miscalibration, with *intensity* exhibiting a systematic underestimation of the gain, and conversely *contrast* and *limbic* exhibiting a systematic overestimation of the gain. Overall, the MSE also tends to be inflated at the extreme of the set sizes considered, when comparing observed and predicted ratio for very small set (*K* =5) and very large sets (*K* =200) (**Fig. S4**). Those deviations suggest the expected gain of multitrait analysis might be refined by including additional features in the model, adding either new parameters, or interactions between those already considered.

**Figure 2.**
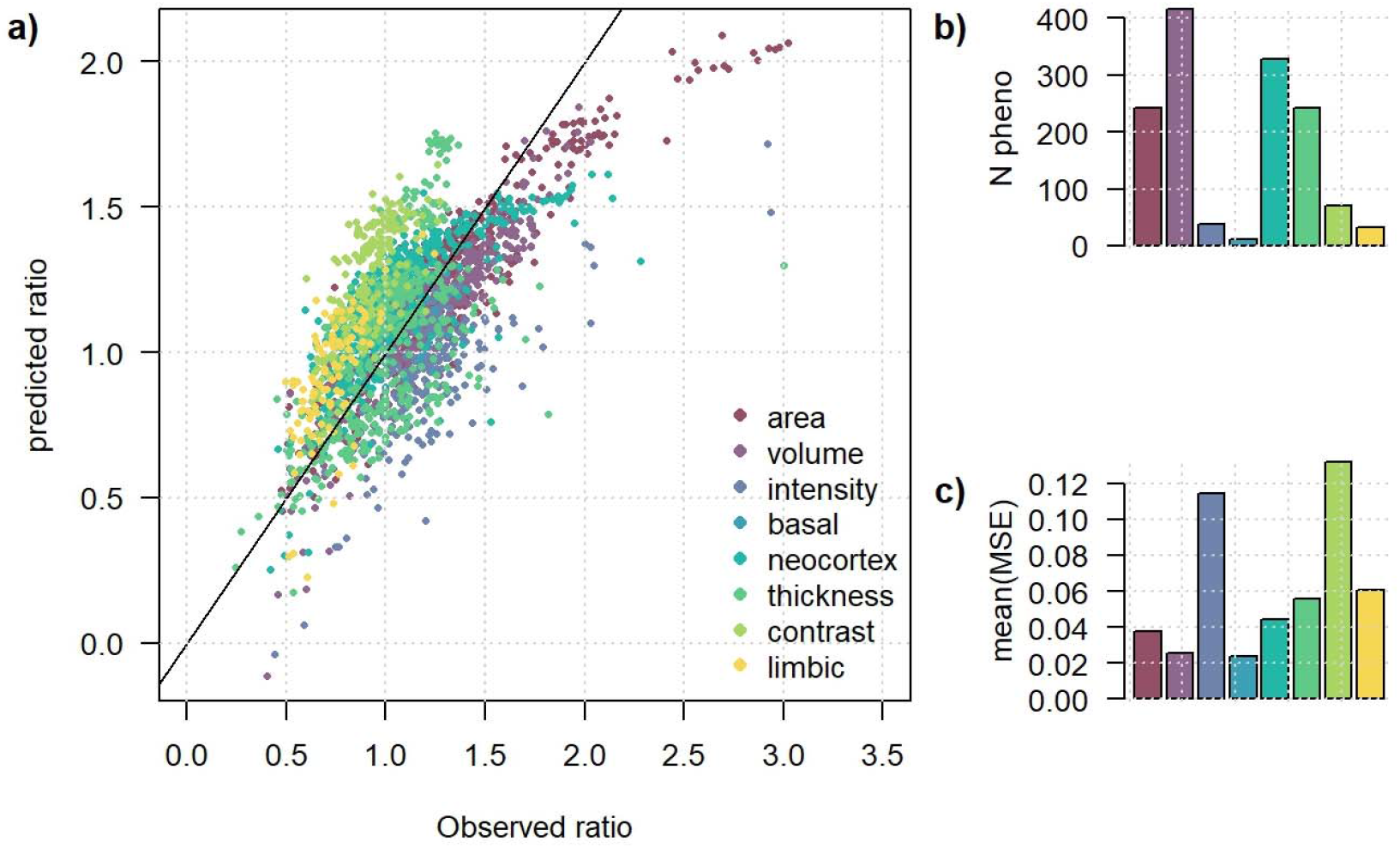
Observed and predicted gain within knowledge-based groups. Observed (X axis) and predicted (Y axis) gain ratio for multitrait against univariate test across eight knowledge-based phenotype groups: area, volume, intensity, basal, neocortex, thickness, contrast, limbic. **(a)** For each group, sets of size 5 to 200 where randomly sampled (with a maximum set size smaller for groups with less than 200 phenotypes) and the observed and predicted gain ratio were plotted against each other’s. **(b)** The groups varied in their characteristics including the number of phenotypes within each of them. **(c)** Although the predicted and observed ratio show high correlation, some groups display a slope offset, producing inflated mean squared error (MSE).

Even though we demonstrated that the gain from multitrait GWAS can be predicted with high accuracy, defining a data-driven approach to determine optimize sets is not trivial. First, we derived our predictive model by sampling sets from the entire dataset, so using it to build optimized sets raise the question of overfitting. However, as suggested in the simulation study from **Figures S5-6** and **Table S5-6**, predictive models build from random sets do not appear to overfit the prediction at other randomly selected sets. In other words, one can train a predictive model by sampling sets from the phenotype pool for which we want to derive optimized sets, which may be intuitively explained by the unsignificant dimension of the training data (∼1000 random sets) as compared to the sampling space (>10^300^ possible sets). Second, this sampling space implies a systematic derivation of the predicted gain for all possible set to select the best sets is obviously unbearable. However, we show that computationally tractable heuristic can be implemented. Here we devised a stepwise procedure were, for a given group of traits of interest, traits are removed iteratively one-by-one until a maximum predicted gain is achieved. We applied this strategy in randomly selected subsets of phenotypes. As showed in **Figure 3**, the observed gain from these optimized sets significantly outperforms the original random sets from **Figure 1**.

**Figure 3.**
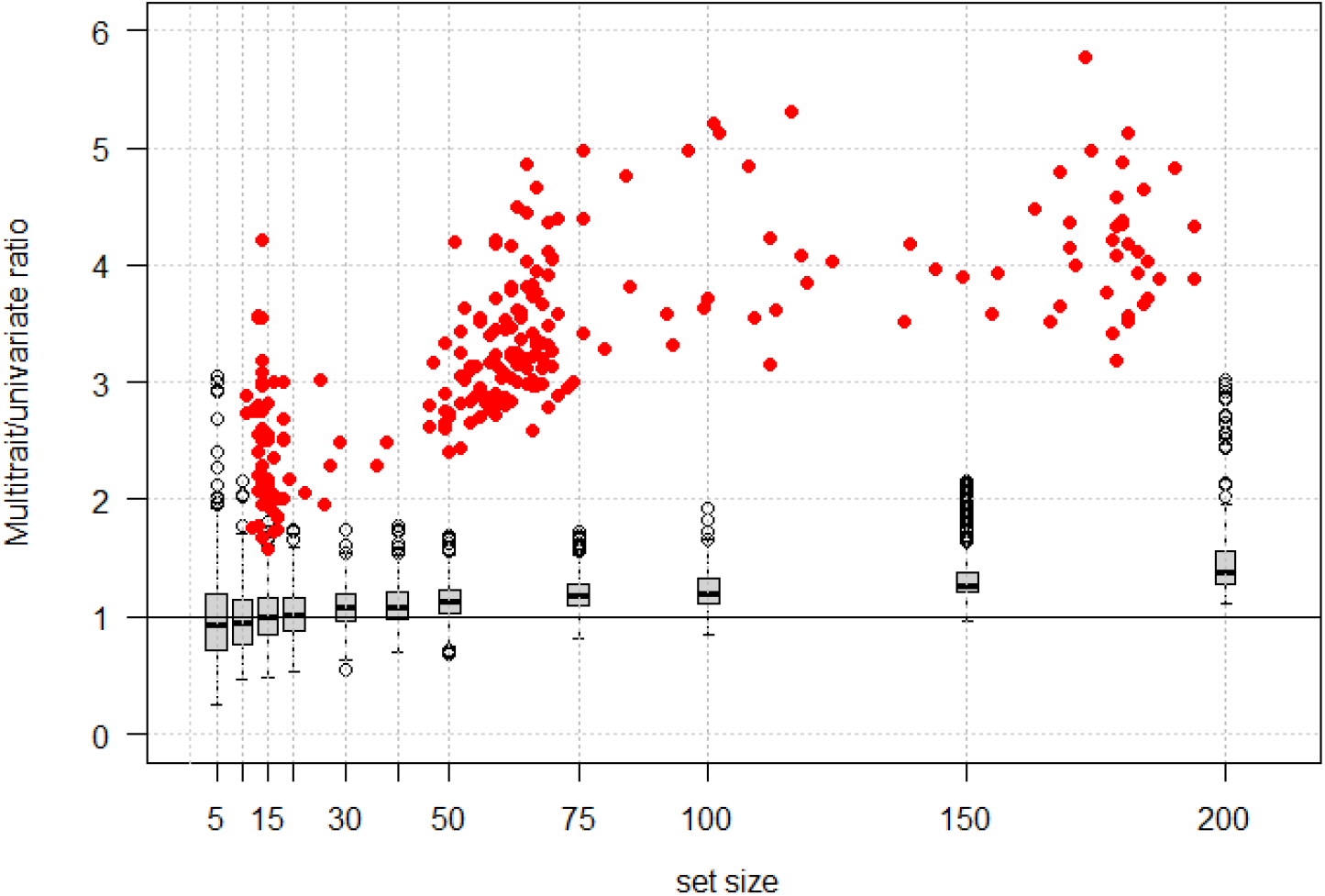
Optimized sets gains compared to random sets. Distribution of the observed multitrait power gain, *τ*_*gain*_, defined as the ratio of independent loci detected by the multitrait test as compared to the univariate test, as a function of set size. Grey boxplots correspond to randomly selected sets. Red dots correspond to sets defined based on the proposed optimized approach, where traits are selected using a stepwise procedure. Given an initial pool of traits, traits are removed one-by-one based on the predicted *τ*_*gain*_. The set with the highest predicted gain is retained and the real gain is derived. The selection was applied for 250 random pools of 50, 100, 125, 175 and 200 traits.

Multitrait GWAS of MRI phenotypes is increasingly used to identify variants missed by standard univariate analysis. Numerous methods have been proposed inside^12,16^ and outside^14,15^ the field of neuroimaging. However, the strategy to select which phenotypes should be tested jointly has been noticeably overlooked. This study demonstrates that the potential power gain of multitrait MRI GWAS over univariate GWAS can be highly predictable, so that in theory, one can define optimized sets to maximize discoveries. Our simulation further suggests that, thanks to the almost unlimited size of the set’s combinatorial space, one can build those predictive models within the data where the multitrait is to be performed without overfitting, thus circumventing the challenge of transferring a predictive model from other independent datasets. Building on this, we show that simple heuristic can be implemented to define optimized set. Optimizing multivariate analysis is not only of practical utility to maximize the use of available data, but can ultimately help mapping the shared genetics between brain MRI and psychiatric disorders^17^.

## Online Methods

### Neuroanatomical brain phenotypes quality control and univariate GWAS

We extracted 1,014 brain phenotypes from UK Biobank for 33,006 unrelated European participants and 9,583,000 variants. The phenotypes include whole brain/left and right hemisphere phenotypes: volumes, area, thickness, grey/white matter contrast, mean intensity and number of holes. For each phenotype *i*, we removed outlier values if they are above or below *threshold*_*i*_ = *mean*(*phenotype*_*i*_) ±3 × *σ* (*phenotype*_*i*_). Four phenotypes with more than 50% missing values were also removed from the analysis: mean intensity and volume of the 5th ventricle / mean intensity and volume of non-white matter hypointensities (all four being whole brain measurements). Phenotypes description (mean, standard deviation and proportion of missing data) used for the analysis are listed in **Table S1**. Univariate GWAS analyses was performed for each phenotype using Plink2^18^, adjusting for sex, age, the top principal components of the genotype matrix, and the UK Biobank assessment center.

### Multitrait *GWAS* and comparison with univariate *GWAS*

We applied an omnibus test implemented in the JASS software^14^, and defined as follows: for each genetic variant, a joint statistic *T*_*omni*_ is derived as: *T*_*omni*_ = *z*^*t*^ *∑*^-1^*z*, where *z* = (*z*_1_, *z*_2_, …, *z*_*k*_) is a vector of *k* Z-scores (for *k* GWAS results), 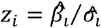. where 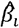. and 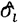. are the estimated regression coefficient and its standard error for trait *i*; and *∑* is the phenotypic correlation matrix between the Z-scores under the null. The latter parameter can be derived from the LDSC^19^ or from individual-level data when available. Here we derived both and confirmed the two estimates are almost fully equivalent (**Fig. S1a** and **Supplementary Material**).

High correlation between variables in linear models can dramatically increase the variance of the estimated effect, which may ultimately impact their robustness. Given the strength of correlation between some of the MRI phenotypes (**Fig. S2**), we devised an *ad hoc* constraint when pulling sets from the whole group to ensure the multitrait test remains calibrated. Briefly, we examined the presence of outliers in the number of detected loci by the multitrait test as a function of various parameters. We devised the following empirical constraint: the set correlation matrix’s condition number had to be smaller than (1.5 x set size) (**Fig. S1b**).

The relative power advantage of the multitrait GWAS over univariate GWAS was computed as *τ*_*gain*_ = *hits*_*multitrait*_/*hits*_*univariate*_ with *hits*_*multitrait*_ and *hits*_*univariate*_ the number of independent loci harboring at least one genome-wide significant variant in the multitrait analysis and in the univariate GWAS, respectively. Independent loci were defined using the 1,703 independent regions previously proposed by Berisa & Pickrell^20^. For the multitrait test we used the standard genome-wide significance threshold of 5 × 10^−8^. For the univariate test, we used a Bonferroni corrected threshold dividing the standard genome-wide significance threshold by the number of traits in the set.

### Parameters associated with multitrait GWAS gain

We evaluated the effect of seven parameters derived for each trait set on the relative power advantage of the multitrait GWAS over univariate GWAS, as measured by *τ*_*gain*_: i) the set size, ii) the average heritability of traits, iii) the mean absolute value of pairwise genetic correlation derived using LDSC^19^, iv) the mean absolute value of the pairwise phenotypic correlation, v) the mean phenotype polygenicity, derived using GCTB^21^, vi) the condition number of the phenotypic correlation matrix, derived as 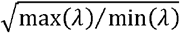 where *λ* is the vector of eigen-values of that matrix, and vii) the number of genome-wide significant hits from the univariate analyses.

The marginal and joint effect of each parameter were estimated using standard linear regression: *τ*_*gain*_ ∼*β*_*i*_ × *Metric*_*i*_, and *τ*_*gain*_ ∼ ∑_*i*_(*β*_*i*_ × *Metric*_*i*_). The latter was also applied within strata defined by the set size to account for potential non-linear effect between set size and the remaining six parameters. Given that the sets used in estimating those coefficients may not be independent because of trait overlap across sets, we also considered a linear mixed model (LMM) modelling this overlap. The LMM was defined as follows: *τ*_*gain*_ ∼*M*^*t*^ Ω with *M* a *q* × *m* matrix for *m* parameters and *q* sets, and Ω a *q* × *q* weighted overlap matrix. The overlap between any pair of two sets *i* and *j* was derived as: *overlap*_*ij*_ = #(*i* ∩ *j*)/#(*i* ∪ *j*). The element of Ω where derived as weighted version defined as 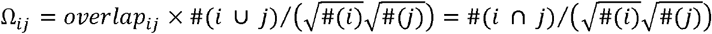.

We used the estimated coefficient of the joint model to derive the predicted gain detection ratio of sets of phenotypes, derived as the sum of the parameters weighted by the coefficients: 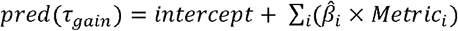.

### Phenotype groups attribution and multitrait analysis

We created 9 groups of phenotypes for the rest of the analysis : one group with all phenotypes (« full data » ; 1,008 phenotypes), five groups based on the phenotypes we selected (« volume », « area », « mean intensity », « G/W matter contrast » and « mean thickness » ; 415, 242, 39, 70 and 242 phenotypes) and three groups based on biological systems, i.e. the basal ganglia with putamen, caudate and pallidum related phenotypes (12 phenotypes), the limbic system with amygdala and hippocampus phenotypes (32 phenotypes) and the neocortex with temporal, occipital, parietal, frontal and thalamus phenotypes (327 phenotypes). For each group, we performed a multitrait analysis on 50 randomly selected subsets of 5, 10, 15, 20, 30, 40, 50, 75, 100, 150, and 200 phenotypes, contingent upon the availability of a sufficient number of traits.

### Optimized selection of phenotypes

We devised a naïve stepwise procedure that leverages the predictive model of the gain to identify optimal multitrait sets. The procedure is defined as follows. For a given initial set of N traits, we calculate the predicted gain of each *N* − 1 combination as *intercept* + ∑_*i*_(*β*_*i*_ × *Metric*_*i*_) and remove the trait that resulted in the maximum predicted gain when removed. The operation is repeated until there are only 5 phenotypes left. The optimal set, displaying the highest estimated gain over all sets considered, is selected. To be implemented, one need the coefficient from the predictive model for each single set size, which we approximated based on the coefficient from the 11 set size used in **Figure 1** (5, 10, 15, 20, 30, 40, 50, 75, 100, 150, and 200) (**Fig. S7a**). We evaluated the performance of this approach in real data. We selected 50 random sets of traits with set size in 50, 100, 125, 175, and 200 and searched for the highest predicted gain while removing phenotypes one by one (**Fig. S7b**). The true gain from of the best sets was then derived based on the real Z-score statistics.

## References

1. Johnstone, EveC., Frith, C. D., Crow, T. J., Husband, J. & Kreel, L. CEREBRAL VENTRICULAR SIZE AND COGNITIVE IMPAIRMENT IN CHRONIC SCHIZOPHRENIA. The Lancet 308, 924–926 (1976).

2. Beutel, M. E., Stern, E. & Silbersweig, D. A. The emerging dialogue between psychoanalysis and neuroscience: neuroimaging perspectives. J. Am. Psychoanal. Assoc. 51, 773–801 (2003).

3. Linden, D. E. J. How psychotherapy changes the brain – the contribution of functional neuroimaging. Mol. Psychiatry 11, 528–538 (2006).

4. Roffman, J. L., Marci, C. D., Glick, D. M., Dougherty, D. D. & Rauch, S. L. Neuroimaging and the functional neuroanatomy of psychotherapy. Psychol. Med. 35, 1385–1398 (2005).

5. Evans, A. C., Janke, A. L., Collins, D. L. & Baillet, S. Brain templates and atlases. NeuroImage 62, 911–922 (2012).

6. Thompson, P. M. et al. The ENIGMA Consortium: large-scale collaborative analyses of neuroimaging and genetic data. Brain Imaging Behav. 8, 153–182 (2014).

7. Zhao, B. et al. Genome-wide association analysis of 19,629 individuals identifies variants influencing regional brain volumes and refines their genetic co-architecture with cognitive and mental health traits. Nat. Genet. 51, 1637–1644 (2019).

8. Elliott, L. T. et al. Genome-wide association studies of brain imaging phenotypes in UK Biobank. Nature 562, 210–216 (2018).

9. Grasby, K. L. et al. The genetic architecture of the human cerebral cortex. Science 367, eaay6690 (2020).

10. Wu, C. Multi-trait Genome-Wide Analyses of the Brain Imaging Phenotypes in UK Biobank. Genetics 215, 947–958 (2020).

11. Sha, Z., Schijven, D., Fisher, S. E. & Francks, C. Genetic architecture of the white matter connectome of the human brain. Sci. Adv. 9, eadd2870 (2023).

12. Tissink, E. P. et al. Abundant pleiotropy across neuroimaging modalities identified through a multivariate genome-wide association study. Nat. Commun. 15, 2655 (2024).

13. Suzuki, Y. et al. Trait selection strategy in multi-trait GWAS: Boosting SNPs discoverability. BioRxiv Prepr. Serv. Biol. 2023.10.27.564319 (2023) doi:10.1101/2023.10.27.564319.

14. Julienne, H. et al. JASS: command line and web interface for the joint analysis of GWAS results. NAR Genomics Bioinforma. 2, lqaa003 (2020).

15. Julienne, H. et al. Multitrait GWAS to connect disease variants and biological mechanisms. PLOS Genet. 17, e1009713 (2021).

16. Shadrin, A. A. et al. Vertex-wise multivariate genome-wide association study identifies 780 unique genetic loci associated with cortical morphology. NeuroImage 244, 118603 (2021).

17. Auvergne, A. et al. Multitrait analysis to decipher the intertwined genetic architecture of neuroanatomical phenotypes and psychiatric disorders. Biol. Psychiatry Cogn. Neurosci. Neuroimaging S2451-9022(24)00266–0 (2024) doi:10.1016/j.bpsc.2024.08.018.

18. Chang, C. C. et al. Second-generation PLINK: rising to the challenge of larger and richer datasets. GigaScience 4, 7 (2015).

19. Bulik-Sullivan, B. et al. An atlas of genetic correlations across human diseases and traits. Nat. Genet. 47, 1236–1241 (2015).

20. Berisa, T. & Pickrell, J. K. Approximately independent linkage disequilibrium blocks in human populations. Bioinformatics 32, 283–285 (2016).

21. Lloyd-Jones, L. R. et al. Improved polygenic prediction by Bayesian multiple regression on summary statistics. Nat. Commun. 10, 5086 (2019).

