## Supplementary data for "Empowering multitrait brain phenotypes GWAS"

**Supplementary Material**

**Table of content**

### **Supplementary** Note

#### Distribution of the correlation between individual GWAS summary statistics under the null

The expected correlation between GWAS z-score summary statistics from two traits $i$ and $j$ equals $cor\left( z_{i},z_{j} \right)={\rho_{ij}N_{s}}/{\sqrt{N_{i}N_{j}}}$, where $\rho_{ij}$ is the phenotypic residual correlation (after adjusting for the covariates included in the GWAS) between traits $i$ and $j$, $N_{i}$ and $N_{j}$ the sample of GWAS of traits $i$ and $j$, and $N_{s}$ the sample overlap. In the special case of the present study where individual-level data is available and the sample overlap is complete, so that $N_{i}=N_{j}=N_{S}$, the summary statistics correlation matrix can be directly derived from the phenotype as $cor\left( z_{i},z_{j} \right)= \rho_{ij}$. When individual-level data are unavailable or the sample overlap unknown, the correlation is typically estimated using the LDSC approach^1^. In the present analysis we used correlation derived from individual-level data. However, as a sensitivity analysis, correlation was also derived using LDSC. As showed in **Figure S2**, the two approaches produce the same estimates.

#### Multivariate GWAS analysis computational cost

There exist a few tools to conduct multitrait analysis from individual-level data, however, most have not been updated to the latest speed-up. For example, a multitrait analysis using three traits, a cohort of 1,000 individuals and 1,000 variants can be done in on average 23 seconds using 2 cores and 24 GB ram using mv-PLINK^2^. mv-PLINK is based on PLINK 1.06 and, as of today, has been implemented neither in PLINK 1.9 nor in PLINK 2.0 which are estimated to be hundreds of times faster than PLINK 1.9 (<https://www.cog-genomics.org/plink/2.0/assoc>). Although updated multivariate software may be released, the computational cost would remain tremendous. The computation cost for standard fixed effect linear regression has $O(N)$ complexity. For large $N$, the standard multitrait analysis can be approximated to $O(NK)$ where $K$ is the number of traits analyzed jointly. It follows that running $M$ multivariate test with an average of $K$ trait would be $O(NKM)$. The average computation time for the univariate GWAS conducted in the UKB dataset with PLINK 2.0, was 10 minutes using 22 CPU (1 CPU per chromosome) and 2.5Go per CPU, or $\sim3$ CPU hours. In our primary analysis, we conducted over 2,500 multivariate GWAS with an average of 50 traits per set corresponding to a total of 2,500 * 50 * 3 = 375,000 hours of computation on a single CPU. In comparison, the same multivariate analysis derived from univariate GWAS summary statistics were derived in in few hours.

#### Study of the overfitting risk

We investigated whether using random sets of phenotypes to train a predictive model of an outcome of interest (here the power gain of the multitrait test) and using the estimated parameters to select optimize sets in the same data would result in overfitting. We used the six real parameters measured in the 2,750 random sets and for each set size we considered in previous analyses (N=[5,…200]), we generated series of 500 simulated null variables, representing an outcome $\Phi$ of interest. For each iteration, we selected 400 new sets of phenotypes. The overlap between set was accounted for by modelling $\Phi$ as a multivariate normal distribution $\Phi\sim N(1,\Omega)$, where the covariance between two set $i$ and $j$ equal $\Omega_{ij}={\#(i \cap j)}/\left( \sqrt{\#(i)}\sqrt{\#(j)} \right)$, as defined in the linear mixed model described in the **Online Methods**. For each replicate we randomly selected 200 sets to train a predictive model of $\Phi$, and tested the association between the predicted $\Phi$ and its simulated value in the remaining sets. To assess the impact of the overfitting, we compared the significance of the adjusted R² in the train data with the *P*-value of the linear regressions between the observed and predicted $\Phi$. As showed in **Figure S5** and **Table S5**, we did not observe any correlation between the two metrics, suggesting that the overlap of traits across sets between training and testing did not resulted in overfitting. Using some of the exact same sets in training and testing on the other hand did produced some overfitting (**Table S6**, **Figure S6**). However it would require a substantial amount of identical sets to encounter an overfitting which should not happen considering the almost infinite number of combinations (>10^300^ for 1,000 phenotypes) in this dataset.

### **ssSupplementary** Figures

#### Figure S1. Multivariate test parametrization and set constraining

Distribution of the phenotypic correlation matrix’s condition number, derived as $\sqrt{{\max\left( \lambda\right)}/{\min\left( \lambda\right)}}$ where $\lambda$ is the vector of eigen-values of that matrix, plotted as a function of the set size. All sets displaying a correlation matrix’s condition number larger smaller than 1.5 times the set size (the red dash line) were filtered out from the analysis.

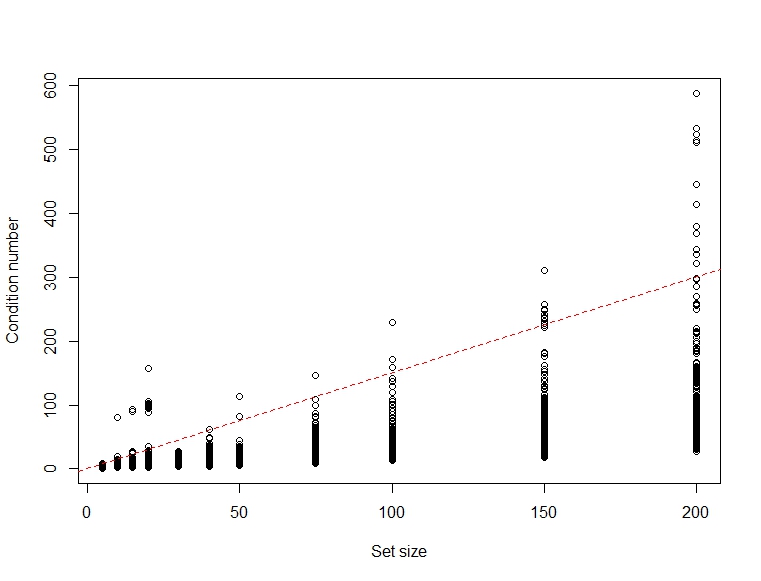

#### Figure S2. Phenotypic and genetic correlation

We derived the phenotypic correlation derived using phenotypes adjusted for the covariates used in the GWAS **(a)** and the genetic correlation estimated using the LDSC approach **(b)** across all 1,010 phenotypes. Panel **(c)** presents the two metrics against each other for all 509,545 pairs of phenotypes. Panel **(d)** compare the expected correlation under the null between GWAS summary statistics across pairs of traits using the LDSC software (Y axis) and the correlation derived from the individual-level data (X axis).

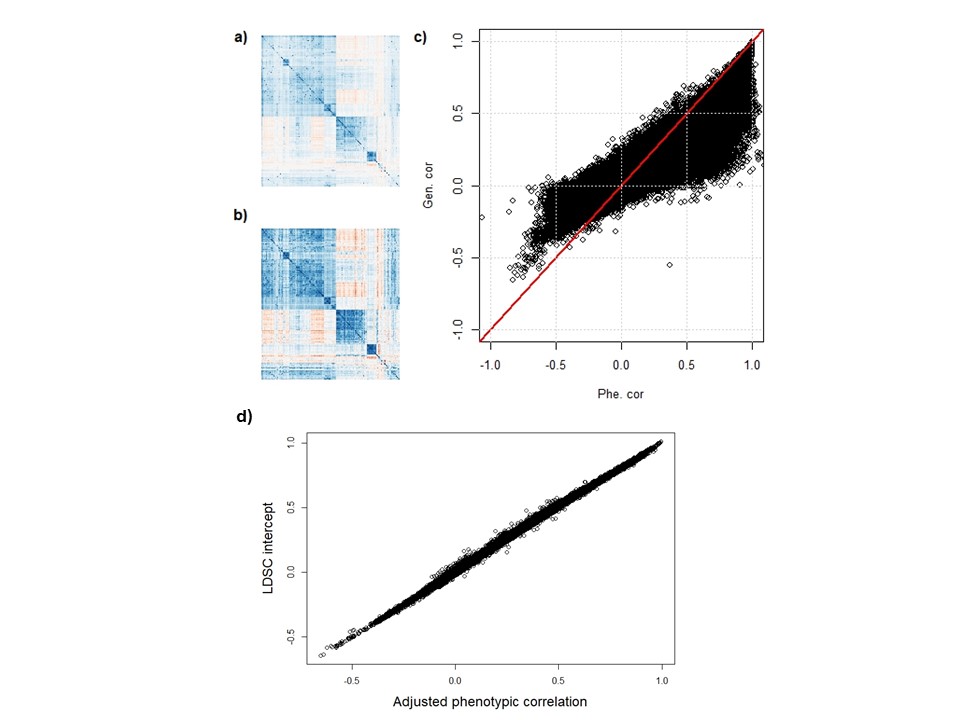

#### Figure S3. Deviation of the estimated gain compared to the observed gain

Observed (X axis) and predicted (Y axis) gain ratio for multitrait against univariate test across eight knowledge-based phenotype groups: area, volume, intensity, basal, neocortex, thickness, contrast, limbic. For each group, sets of size 5 to 200 where randomly sampled (with a maximum set size smaller for groups with less than 200 phenotypes) and the observed and predicted gain ratio were plotted against each other’s.

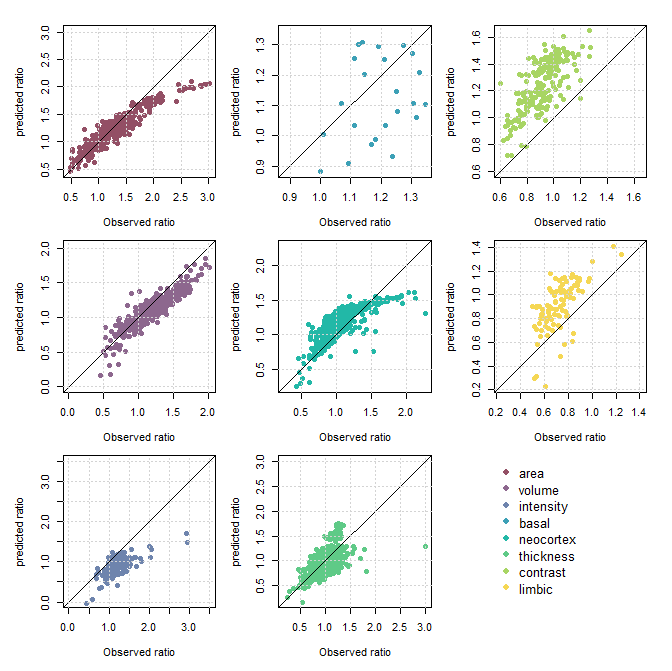

#### Figure S4. Deviation of the estimated gain per size set

Difference between the observed and predicted gain for multitrait against univariate test as measured by the mean squared error (MSE) between the observed and predicted ratio as a function of the set size. Results are presented for each of eight knowledge-based phenotype groups considered: area, volume, intensity, basal, neocortex, thickness, contrast, limbic. The plain red lines indicate the average MSE within each set size.

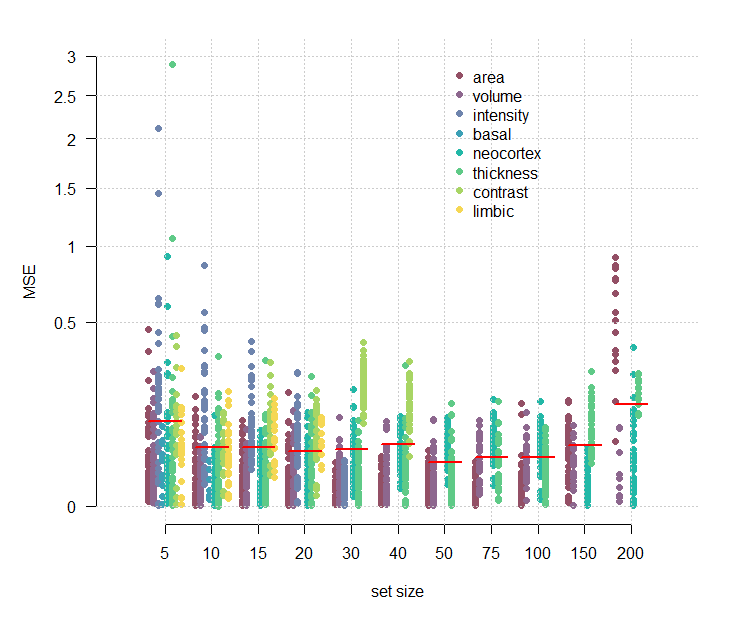

#### Figure S5. Evaluation of overfitting

We assessed potential overfitting issues of training a predictive model of the multitrait gain in the same data where the gain is to be predicted. We used the real six parameters measured in the 2,750 random sets from Figure 1a and generated replicates of 500 sets were a null variable mimicking multitrait power gain was simulated. Each replicate was split into training and test. The train data were used to build a predictive model of the null variable and the R² of that model was recorded. The predictive model was then applied in the test data to predict the null variable, and the association between real and predicted null variable was derived using a standard linear regression. Correlation between the train R² *p*-value (X axis) and the predication accuracy in the test data (Y axis) was derived as a measure of overfitting. The simulation was repeated for set sizes in [5, 200]. The red line display $x=y$.

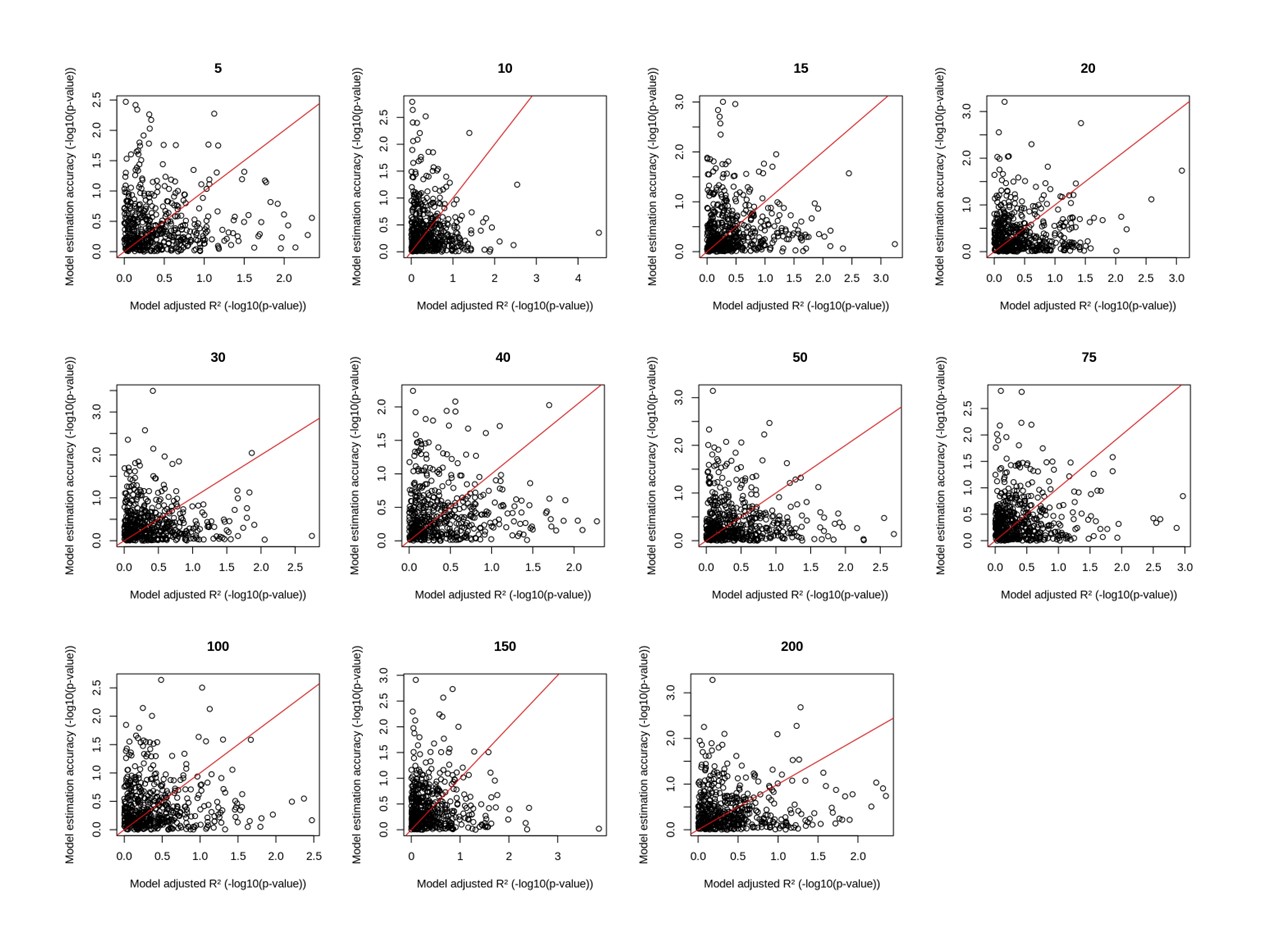

#### Figure S6. Overfitting in the special case of strong overlap

We assessed potential overfitting issues of training a predictive model of the multitrait gain in the same data where the gain is to be predicted in the special case where some exact same sets are part of the training and testing data. We used the real six parameters measured in the 2,750 random sets from Figure 1a and generated replicates of 500 sets were a null variable mimicking multitrait power gain was simulated. Each replicate was split into training and test, although a portion of the training sets were also used in the test set. The train data were used to build a predictive model of the null variable and the R² of that model was recorded. The predictive model was then applied in the test data to predict the null variable, and the association between real and predicted null variable was derived using a standard linear regression. Correlation between the train R² *p*-value (X axis) and the predication accuracy in the test data (Y axis) was derived as a measure of overfitting. The simulation was repeated for set sizes in [5, 200]. The red line display $x=y$.

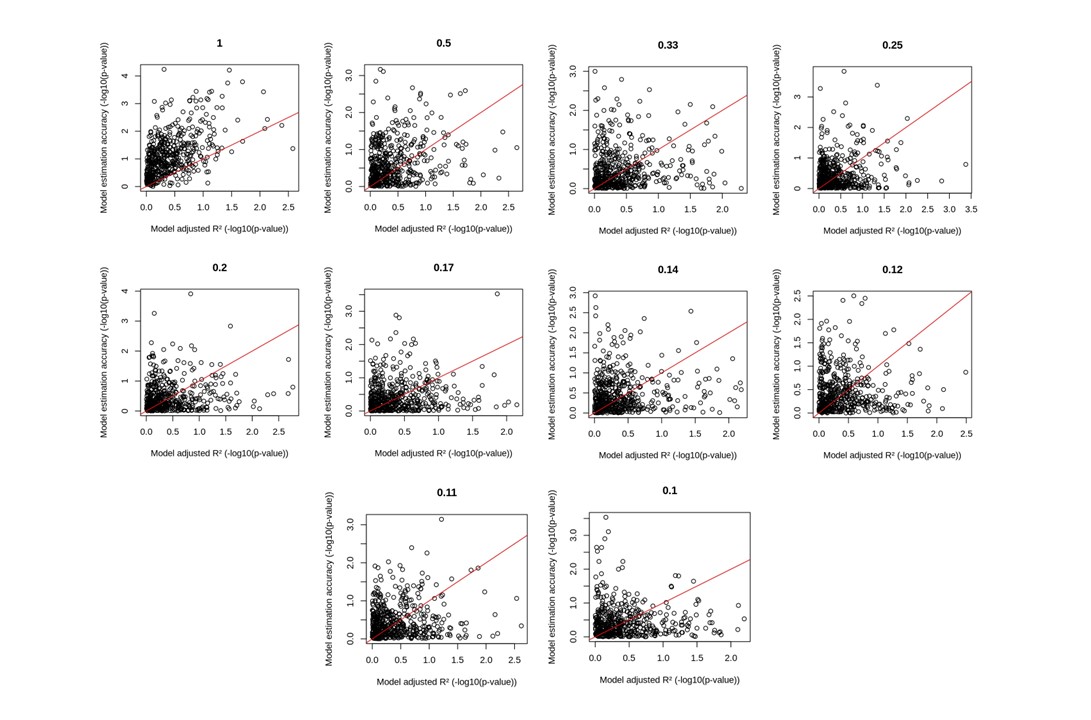

#### Figure S7. Optimized selection of phenotypes

We derived a continuous prediction model by approximating the coefficients from the six parameters for set sizes varying from 5 to 250. Panel a) shows the estimated parameters’ coefficient (black dots), with their size representing their weight for the approximation, and the approximated coefficients (red line). Panel b) displays the boxplots of the predicted gain from the approximated coefficients for random set of traits, with the mean in red. The blue lines represent the average predicted gain over 50 experiment from the proposed stepwise approach. Starting from a given set size (N=50, 100, 125, 175, and 200, blue dots), traits are removed one-by-one so that the predicted gain from the remaining traits is maximized.

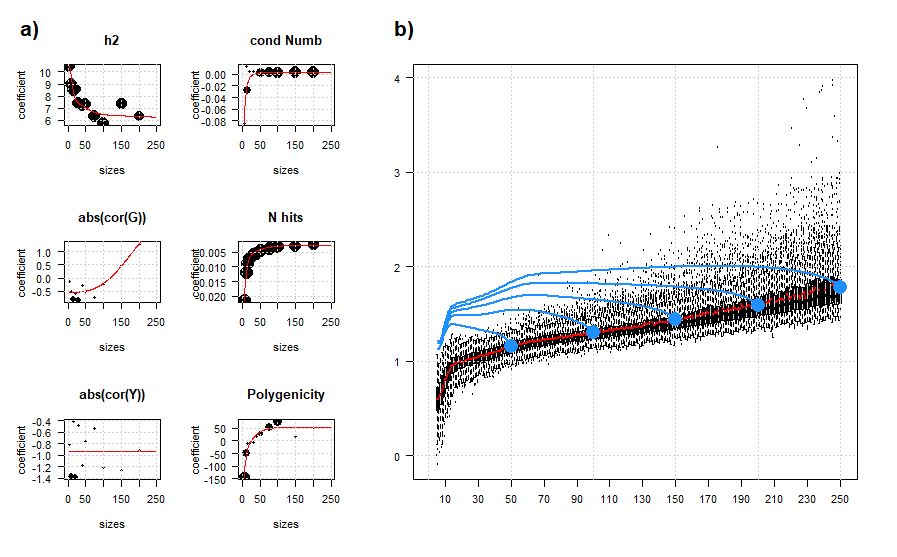
